# Single-Cell Multiomics Reveals Regulatory Mechanisms of CAR T Cell Persistence and Dysfunction in Multiple Myeloma

**DOI:** 10.1101/2025.04.01.646378

**Authors:** Lorea Jordana-Urriza, Guillermo Serrano, Sergio Camara, Maria E. Calleja-Cervantes, Patxi San Martin-Uriz, Aintzane Zabaleta, Aina Oliver-Caldes, Marta Español-Rego, Diego Alignani, Teresa Lozano, Saray Rodriguez-Diaz, Elena Iglesias, Valentin Cabañas, Juan L. Reguera, Veronica Gonzalez-Calle, Maria V. Mateos, Fermin Sanchez-Guijo, Bruno Paiva, Juan J. Lasarte, Susana Inoges, Ascensión Lopez-Diaz de Cerio, Azucena Gonzalez-Navarro, Manel Juan, Carlos Fernandez de Larrea, Esteban Tamariz, Ana Alfonso-Pierola, Paula Rodriguez-Otero, Jesús San-Miguel, Mikel Hernaez, Juan R. Rodriguez-Madoz, Felipe Prosper

**Affiliations:** Hemato-Oncology Program. Cima Universidad de Navarra. IdiSNA. Pamplona, Spain; Computational Biology Program. Cima Universidad de Navarra. IdiSNA. Pamplona, Spain; Department of Hematology. Hospital Clinic de Barcelona. IDIBAPS. Universidad de Barcelona. Barcelona, Spain; Department of Immunology. Hospital Clinic de Barcelona (HCB). Joint Platform HSJD-HCB. IDIBAPS. Universitat de Barcelona. Barcelona, Spain; Immunology and Immunotherapy Program. Cima Universidad de Navarra. IdiSNA. Pamplona, Spain; Department of Hematology, IMIB-Virgen de la Arrixaca University Hospital. University of Murcia. Murcia, Spain; Department of Hematology, University Hospital Virgen del Rocio-IBIS. Universidad de Sevilla. Sevilla, Spain; Hematology Department, IBSAL-University Hospital of Salamanca. University of Salamanca. Salamanca, Spain; Hematology and Cell Therapy Department. Clinica Universidad de Navarra, IdiSNA. Pamplona, Spain; Centro de Investigacion Biomedica en Red de Cancer (CIBERONC). Madrid, Spain; Cancer Center Clinica Universidad de Navarra (CCUN). Pamplona, Spain; Data Science and Artificial Intelligence Institute (DATAI). Universidad de Navarra. Pamplona, Spain

## Abstract

Understanding the mechanisms that drive chimeric antigen receptor (CAR) T cell function and persistence in multiple myeloma (MM) remains a critical challenge for improving therapeutic outcomes. In this study, we applied single-cell multiomics and gene regulatory network (GRN) analysis to characterize the transcriptional dynamics and clonal evolution of BCMA-targeted CAR T cells in longitudinally collected bone marrow (BM) and peripheral blood (PB) samples from MM patients. Our results revealed that CAR T cells infiltrating BM exhibited a more activated and exhausted phenotype compared to their PB counterparts, with key transcriptional regulators driving these changes. Dysregulation in the effector-to-memory transition led to an increased presence of terminally differentiated CAR T cells, correlating with poor persistence. Additionally, we identified a hyperexpanded CAR T clone in the BM of a patient with partial response, marked by elevated IL10 expression. Functional analyses demonstrated that stimulation of endogenous TCR enhanced IL10 production, potentially contributing to impaired CAR T cell proliferation and persistence. These findings uncover critical regulatory mechanisms influencing CAR T cell dynamics, offering new insights into improving CAR T cell persistence and therapeutic efficacy in MM and highlights potential molecular targets for optimizing CAR T cell therapy in patients with MM.

## INTRODUCTION

Chimeric antigen receptor (CAR)-T cell therapies have become a promising therapeutic option for patients suffering B-cell hematological malignancies, including acute lymphoblastic leukemia (ALL), diffuse large B cell lymphoma (DLBCL), and multiple myeloma (MM)^1^. In patients with MM, CAR T therapies targeting BCMA have produced unprecedent rates of remission and increased life expectancy for relapsed/refractory (R/R) patients^2–5^. Nevertheless, not all patients respond to therapy with almost every patient eventually relapsing, with a median PFS of 11-36 months depending months depending on the product^2–5^. This lack of long-term response, one of the main challenges in CAR T therapies for MM, depends on multiple factors, including CAR construct design, the quality and composition of the CAR T cell product, the CAR T cell expansion and persistence capability, the development of resistance mechanisms and/or the tumor burden and tumor microenvironment^6–14^. All these data clearly indicate the pressing need to better understand the underlying cellular and molecular features of CAR T cell therapy leading to different patient outcomes. However, there is still missing knowledge in the factors that may be crucial for a prolonged CAR T cell response in MM, and their identification remains challenging.

The advancement of omic technologies, and particularly those with single cell resolution, has allowed gaining insights in the heterogeneity of CAR T cell products as well as identifying molecular factors driving CAR T function *in vivo*. Moreover, the analysis of Gene Regulatory Network (GRN) has demonstrated to be a powerful tool to unravel regulatory dynamics of CAR T cells^15,16^. However, most of the studies have been conducted analyzing CD19 CAR T cells isolated from the infusion product, or circulating in blood, from patients with ALL and DLBCL^17–22^. Even though there is increasing literature on the transcriptional features driving CAR T cell dynamics in MM^23–25^, there is a still an incomplete knowledge on the molecular factors important for long-term CAR T cell functionality. Moreover, we lack a full understanding of the differences between CAR T cells infiltrating the bone marrow (BM) and their counterparts circulating in peripheral blood (PB), a critical aspect in MM, due to the nature of the disease. Thus, there is a critical need to study the dynamics and evolution of CAR T cell after infusion in both compartments to increase our knowledge and identify key factors responsible of long-lasting CAR T cell responses in MM.

In this study we interrogated longitudinal samples of CAR T cells collected from patients presenting different clinical characteristics and outcomes, enrolled in the academic clinical trial CARTBCMA- HCB-01 (NCT04309981)^26,27^, assessing the efficacy and safety of BCMA CAR T. We applied single cell RNA (scRNA-seq) and TCR sequencing (scTCR-seq) coupled with SimiC^15^, a machine learning algorithm that infers GRNs, to characterized almost 50.000 CAR T cells from 11 different samples, including infusion products (IP), as well as CAR T cells isolated from BM and PB, at one and three months after infusion. Our data reveal relevant differences in regulatory networks between CAR T cells infiltrating the BM and those in PB, identifying specific regulons as key drivers of the more activated and exhausted phenotype observed in BM CAR T cells. Moreover, we define regulatory dysfunctions in the effector-to-memory axis promoting the lack of CAR T cell persistence. Finally, we identify a hyperexpanded CAR T clone in the BM of a patient with partial response, presenting high expression of IL10, which was associated to expression of transcription factors related to exhausted CAR T cells, providing a potential mechanism for CAR T cell failure in this patient. Our analyses provide insights into the regulatory mechanisms that could represent potential targets to be modulated for the development of improved CAR T therapies for MM.

## MATERIALS AND METHODS

### Patient samples and clinical data

Peripheral blood (PB) and bone marrow (BM) samples were collected at the indicated time points from patients with MM enrolled in the academic clinical trial CARTBCMA-HCB-01, assessing the efficacy and safety of BCMA-CAR T ARI0002h (NCT04309981)^26,27^. All subjects provided written informed consent. Clinical response was defined according to the MM Response Criteria^28^ as: Stringent Complete Response (sCR), Very Good Partial Response (VGPR), Partial Response (PR) and Progressive Disease (PD).

### Flow cytometry and human CAR T cell isolation

IP, PB and BM samples were processed as previously described^16,26^. Briefly, after bulk lysis for erythrocyte elimination, CAR T cells were incubated with scFv-BT (Jackson Immunoresearch), washed with PBS/BSA and stained with Streptavidine-BV421 (BD Biosciences). CAR T cells were sorted either with BD FACSAria IIu or Moflo ASTRIOS BC. Data was acquired on a BD FACSCanto II (BD Biosciences) and analyzed using the FlowJo Software version 10 (Tree Star).

### Single cell RNA-sequencing and single-cell V(D)J sequencing data

FACS-sorted CAR T cells from BM and PB samples of MM patients were subjected to single cell RNA and TCR sequencing (scRNA-seq and scTCR-seq) using the Chromium Single Cell 5′ Reagent Kit (10X Genomics) according to manufacturer’s instructions. After quality control and quantification, single cell libraries were sequenced at a minimum depth of 30000 reads/cell. From 66520 analyzed cells, 47856 passed the quality control yielding an average of 2257 genes/cell. TCRα/β sequencing was performed with 10X Genomics Single Cell V(D)J Immune Profiling Solution (10x Genomics). After quality control and quantification, single cell V(D)J enriched libraries were pooled and sequenced at a minimum depth of 5000 reads per cell. scRNA-seq data were demultiplexed, aligned to the human reference (GRCh38) and the feature-barcode matrix was quantified using Cell Ranger (v6.0.1) from 10X Genomics. Further computational analysis was performed using Seurat (v3.1.5). Cells with no detectable expression of CD3 or CAR were computationally removed, as well as cells with less than 250 genes and with more than 30000 unique molecular identifiers (UMIs). Moreover, any cells that were predicted to be doublets with Scrublet were removed. Finally, cells with a mitochondrial RNA greater than 10% were also removed. All scRNA-seq count matrices were merged and log-normalized. Batch correction across patients was performed with Harmony using default parameters. Harmony was applied to the principal components derived from the top 2000 variable genes. Dimensionality reduction was performed using t-distributed stochastic neighbor embedding (t-SNE) and unsupervised clustering analysis with the resolution set to 0.8. Clusters were named using canonical marker genes as reference. TCR analysis was performed using scRepertoire package.

### Gene regulatory network analysis

Using the most variable 100 TFs and 1,000 genes, SimiC^15^ was run with the default parameters. For pre- vs post-infusion GRN analysis **λ**1 and **λ**2 were set to 0.001 and 0.01, respectively, and for BM vs PB analysis both **λ** hyperparameters were set to 0.001. For the inference of IL10 regulon, a list of the most variable 300 TFs in post infusion CAR T cells were selected and IL10 as the unique target, with 0.001 and 0.01 values for **λ**1 and **λ**2, respectively. For each regulon, defined as a TF and its associated target genes, activity score per cell is computed as the area under the curve generated by the cumulative sum of the ordered weights corresponding to the target genes connected to the TF. Then, the distribution for each cell phenotype is represented in a histogram plot, and the regulatory dissimilarity of each regulon is calculated computing the distance between the distribution of the cells belonging to each phenotype. GRNs were plotted using the GRN incidence matrices provided by SimiC.

### TCR signature generation

TCR signature was based on the Affymetrix microarray data by Yang et al^29^. Briefly, the transcriptomic data of CD8^+^ T cells stimulated through the TCR and CAR were compared to CD8^+^ T cells stimulated only through the CAR. Then, selected genes were converted to their human orthologs and only those that were expressed by cells in our single cell data object were kept. After filtering steps, the signature comprised a total of 79 genes.

### Cell lines

The Platinum Ecotropic (Plat-E, ATCC) cell line was cultured in DMEM supplemented with 10% FCS at 37°C in a humidified atmosphere with 6.5% CO2. MM5080 cell line^30^ was cultured in RPMI supplemented with 10% FBS at a humidified atmosphere with 6.5% CO2.

### Retroviral vector construction and virus preparation

Retroviral vector (RV) coding the murine CAR construct targeting mouse BCMA was generated as previously described^31^. Briefly, a pRubiG retroviral vector was used to express a murine CAR comprising an anti-mouse BCMA single-chain variable fragment (scFv), a CD8 hinge and transmembrane domain, the 4-1BB and CD3ζ endodomains fused to a GFP reporter gene. PlatE cells were transfected with 5μg of retroviral plasmid and 2.5 μg pCL-Ecoplasmid DNA using Lipofectamine 2000 (Invitrogen) for 6 hours in antibiotic- free medium. Supernatants were collected at 48h and 72h after transfection.

### Murine CAR T cell generation

Spleens and lymph nodes of CD45.1 OT-1 mice were extracted for the isolation of CD8^+^ T cells. Single-cell suspensions were obtained by homogenizing the organs with a cell strainer in ACK lysis buffer. Then, CD8^+^ cells were magnetically separated with the CD8a^+^ T Cell Isolation Kit (Miltenyi) following manufacturer’s instructions. Purified cells were cultured in complete RPMI medium containing 10% FBS, 1% penicillin/streptomycin, 1% L-Glutamine, 1% NEAA, 1% Na-Piruvate, 0,1% anphotericine B, 0,1% B-mercaptoetanol and 1mM HEPES. Purified mouse CD8^+^ T cells were activated with CD3/CD28 dynabeads at a 1:2 bead:T cell ratio for 24h (10^6^ cells/mL density in 12-well plates) in RPMI complete media containing 100 IU/mL interleukin-2 (IL-2). 24h after activation, CD8^+^ cells were collected and cultured for 48h with retroviral supernatant with 100 IU/mL IL-2 and 10 μg/mL protamine sulfate (Sigma) and centrifuged at 2000xg at 32°C for 90 min in 12 well plates. 24h after the first infection the infective medium was removed, and cells were re-infected with the 72h retroviral supernatants. After infection, lymphocytes were cultured in complete RPMI medium with IL-2 for 24h. BCMA CAR T cells were FACS-purified based on the expression of GFP.

### Analysis of murine CAR T cell response to BCMA, SIINFEKL and IL-10

OT-1 BCMA CAR T cells were seeded at a density of 1x10^5^ cells/well in 96-well plates pre-coated with recombinant mouse BCMA (rBCMA) (R&D Systems) (30 ng/ml). SIINFEKL (0,1ng/ml) was added in the corresponding wells diluted in complete RPMI medium. After 24h of incubation supernatants were collected for cytokine detection. For analysis of CAR TCR sequential stimulation, OT-1 BCMA CAR T cells were first seeded in rBCMA pre-coated plates and after 48h of incubation they were collected, counted and cultured in new plates in complete RPMI medium supplemented with SIINFEKL. After 24h supernatants were collected for cytokine analysis. In other experiments, CAR T cells were incubated in normal complete RPMI medium or supplemented with 5ng/ml mouse recombinant IL10 (rIL10) (PeproTech) for 3 days. Then, 1x10^5^ cells/well were seeded in 96-well plates pre-coated with rBCMA in complete medium containing 5ng/ml rIL10. After 24h CAR T cell proliferation and IFN-γ production were measured.

### Analysis of murine CAR T cell proliferating and cytokine quantification

Proliferation was measured by [methyl-3H]-thymidine incorporation. Briefly, cells were centrifuged at 2000rpm for 5min, the supernatant removed and 0.5 μCi of [methyl-3H]thymidine was added to each well and incubated for 6-8 hours. Cells were harvested (Filtermate 96 harvester; Packard Instrument), and incorporated radioactivity was measured using a scintillation counter (TopCount; Packard Instrument). IL10 and IFN-γ production were quantified using mouse ELISA sets (BD OptEIA), following manufacturer’s instructions. The number of IFN-γ-producing CAR T cells was measured by ELISPOT assay (BD-Biosciences). Briefly, CAR T cells were plated for 24 hours in the presence of irradiated MM5080 cells at different tumor:CAR ratios (1:1, 0,25:1, 0,1:1 and 0:1). After one day of culture the number of spot-forming cells was enumerated with an automated ELISPOT reader (CTL, Aalen, Germany).

### Statistical Analysis

Statistical analyses were performed using R version 4.1.3 and GraphPad Prism version 10.1.0. The different tests used in this work are indicated in the figure legend.

## RESULTS

### Expansion of CAR T cells after infusion is preferentially driven by CD8+ T cells

To deepen into the transcriptomic programs governing CAR T cell dynamics through therapy, we collected longitudinal samples of CAR T cells from MM patients treated with BCMA CAR T therapies enrolled in CARTBCMA-HCB-01 (NCT04309981)^26,27^ clinical trial. Specifically, CAR T cells were collected from infusion products (IPs) as well as paired bone marrow (BM) and peripheral blood (PB) samples, at one and three months after infusion, from three patients with R/R MM (Figure 1A and 1B). Then, FACS-isolated CAR T cells were profiled using single cell RNA (scRNA-seq) and single cell TCR (scTCR-seq) sequencing. After quality control and filtering, a total of 47,855 CAR T cells (expressing CD3 and CAR) (Figure S1A) were included in the analysis (median of 3,847 cells per sample, range 384-10,282 cells) (Table S1). Unsupervised clustering of integrated samples yielded 12 clusters, composed by CAR T cells proceeding from all three patients that were defined according to the expression of canonical markers^16^ (Figure 1C, 1D and S1B). We observed remarkable differences between IP and post-infusion samples, as reported previously^20^, with six clusters belonging almost exclusively to the IP (Figure 1D). These clusters included a diversity of CAR T cell populations with proliferating cells (both CD4^+^ and CD8^+^), activated CD4^+^, memory CD4^+^ and CD8^+^, as well a small population of CD8^+^GNLY^-^ effector cells (Figure 1C, 1D and S1A). On the other hand, post-infusion CAR T cells (additional six clusters) were mainly composed by non-proliferating CD8^+^ CAR T cells with effector and effector-memory phenotypes, except for three groups with small number of cells corresponding to proliferative and resting CD8^+^ cells and early-memory CD4^+^ cells (Figure 1C, 1E and S1A). These results clearly show a preferential expansion of CD8^+^ subset after infusion in these patients, that was also observed in most of the patients enrolled in this trial (Figure S1C), that would be in accordance with the requirement of highly cytotoxic effector cells during the early phases of the CAR T cell therapy.

**Figure 1.**
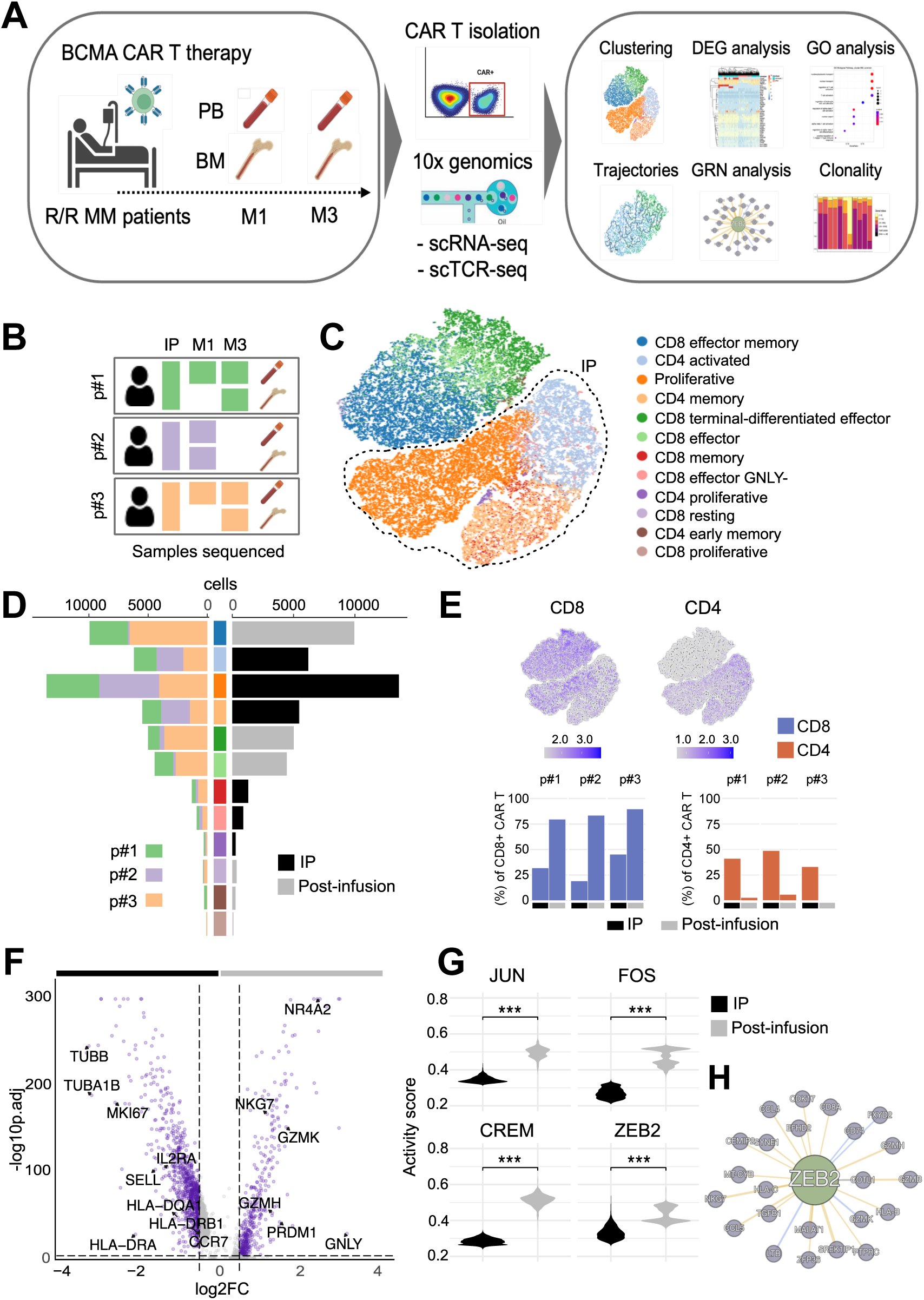
Preferential expansion of CD8^+^ CAR T cells with effector transcriptomic profiles after infusion. Transcriptomic analyses were performed on CAR T cells collected from MM patients at different times after infusion (M1 and M3) and their paired infusion product (IP). **(A)** Schematic representation of the followed pipeline. CAR^+^ cells were FACS-isolated sorted from PB and BM samples. Then, RNA was extracted and scRNA-seq and scTCR-seq were performed. **(B)** Schematic representation of samples analyzed. **(C)** t-SNE plot of the 47,855 CAR T cells that passed quality control and filtering for subsequent analysis in the study. CAR T cell populations were defined according to the expression of canonical markers. **(D)** Relative contribution of each sample to the defined CAR T cell populations in terms of patient (left) or timepoint (right). **(E)** t-SNE plots showing the expression of CD8A and CD4, with the proportions of CD8^+^ and CD4^+^ CAR T cells in IP and post-infusion samples in each patient. **(F)** Differentially expressed genes between CD8^+^ CAR T cells in IP vs post-infusion samples (p adjusted < 0.01 and logFC > |0.25|). **(G)** Activity scores of regulons of interest in IP vs post-infusion samples. **(H)** Representation of ZEB2 regulon depicting target genes with a positive (yellow) or negative (blue) correlation. Wilcoxon test for unpaired samples (G, H) with Boferroni or Benjamini-Hochberg (BH) procedures for p value correction (G and H, respectively). ***p < 0.001.

Differential expression analysis showed that IPs overexpressed factors related to proliferation (MKI67, TUBB, TUBA), activation (HLA class II genes, CD25) and memory (CD62L, CCR7), according to the different CAR T cell populations present on IPs (Figure 1F and Table S2). In contrast, post-infusion CAR T cells presented higher expression of genes related to cytotoxicity, like NKG7, GZMK, GZMH and GNLY, that together with the overexpression of factors related to exhaustion like NR4A2^32,33^ and PRDM1^34^, suggested a more terminally differentiated profile of those expanded CD8^+^ CAR T cells (Figure 1F and Table S2). To have a more comprehensive view on the mechanisms driving CAR T cell function after infusion, we performed a GRN analysis with SimiC^15^. We observed 63 regulons presenting increased activity scores in post-infusion samples compared to IPs (Figure S1D and Table S3). Among the most differentially active regulons we found TFs involved in the downstream signaling of T cells, like JUN and FOS^35^, as well as TFs related to T cell terminal differentiation and exhaustion, like CREM^36^ and ZEB2^37^ (Figure 1G and S1E). Moreover, ZEB2 regulon was positively correlated with T cell effector genes (GZMB, NKG7, CCL5 or GZMM) (Figure 1H). Overall, these results supported the transcriptomic profiles observed in CAR T cells after infusion. In addition, GRN analysis also revealed increased activity of several zinc finger factors in these cells, like ZNF800, ZNF292 and ZNF394, that have been associated to cell differentiation^38^, cell cycle suppression^39^ and AP1/c-JUN inhibition^40^ respectively (Figure S1F). Although the function of these regulons in T cells is still largely unknown, they could represent novel potential regulators of CAR T cell function *in vivo*.

### Post-infusion CD8^+^ CAR T cells exhibit an effector-to-memory phenotypic transition

Next, we performed a deeper characterization of the most relevant CD8^+^ CAR T cell populations observed post-infusion in BM and PB. Within the effector phenotype we observed two transcriptionally distinct populations, denoted as CD8^+^ terminally-differentiated-effector (CD8^+^TE) and CD8^+^ effector (CD8^+^Eff) (Figure 2A). CD8^+^TE expressed higher levels of activation (HLA-DRA, HLA-DRB1, HLA- DQA1, TNFRSF9/4-1BB), cytotoxicity (GZMK, GZMA, IFNG), exhaustion (LAG3, CTLA4, HVACR2/TIM3) and terminal differentiation markers (PRDM1, NR4A2) as revealed by DGE analysis (Figure 2B and Table S4). Moreover, CD8^+^TE scored higher in effector-related hallmark signatures like IFNγ, TNFα signaling and apoptosis (Figure 2C). In contrast, CD8^+^Eff population was enriched in translation and ribosome biogenesis pathways when compared to the CD8^+^TE population (Figure S2A). These CD8^+^TE and CD8^+^Eff cell subpopulations, together with the CD8^+^ effector-memory cells (CD8^+^EM), composed a post-infusion effector-to-memory axis (Figure 2A).

**Figure 2.**
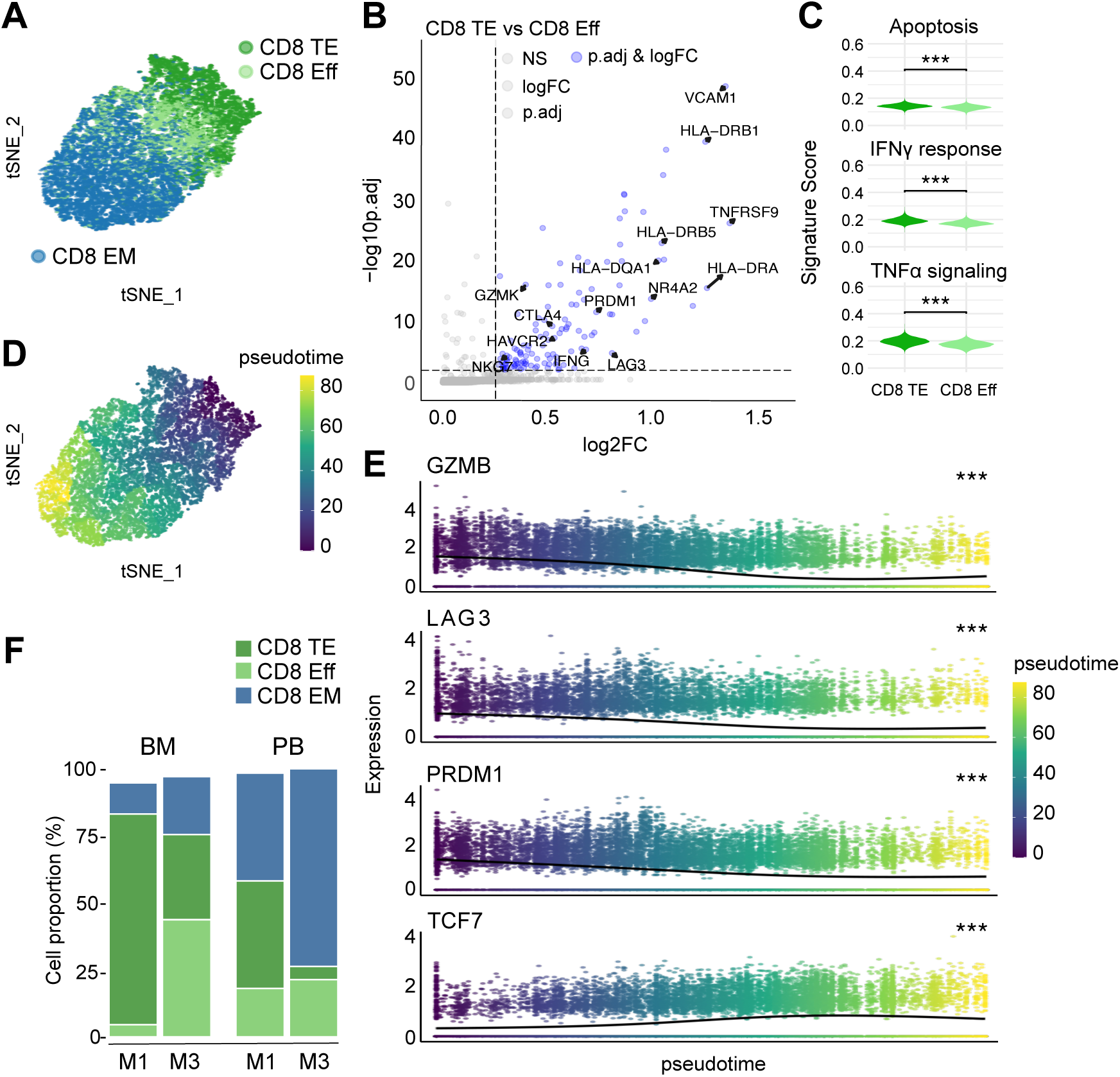
Post-infusion CAR T cells undergo a phenotypic transition along the effector-to-memory axis. Post-infusion CAR T cells were further characterized, focusing on the CD8^+^ cells with effector and effector-memory phenotypes. **(A)** t-SNE plots showing distribution of CD8^+^TE, CD8^+^Eff and CD8^+^EM populations within post-infusion CAR T cells. **(B)** Differentially expressed genes between CD8^+^TE and CD8^+^Eff CAR T cells (p adjusted < 0.01 and logFC > 0.25). **(C)** Scoring for apoptosis, IFNγ and TNFα gene signatures in CD8^+^TE and CD8^+^Eff CAR T cells. **(D)** t-SNE plot showing pseudotime analysis spanning through the effector-memory axis of post-infusion CAR T cells. **(E)** Expression of GZMB, LAG3, PRDM1 and TCF7 genes along the pseudotime. **(F)** Proportion of main post-infusion CAR T clusters in BM and PB at moths 1 and 3 after infusion. Wilcoxon test for unpaired samples (B, C) with Bonferroni or Benjamini-Hochberg (BH) procedures for p value correction (B and C, respectively). Natural cubic splines regression (E). ***p < 0.001.

To better understand the transcriptional dynamics across this effector-to-memory axis, we performed gene trajectory analysis across the pseudotime defined by such axis (Figure 2D). We found that expression of cytotoxicity markers (GZMB, GZMK, LAG3, PRF1) and TFs related to effector commitment (PRDM1, ZEB2) decreased along the pseudotime, with a concomitant upregulation of TFs associated to memory (TCF7, FOXP1) (Figure 2E and S2B). Further, when mapping this phenotypic axis to the course of therapy, CD8^+^TE cells were more abundant at month 1, meanwhile CD8^+^Eff and CD8^+^EM cells were present at longer times (month 3) (Figure 2F), indicating that CAR T cells follow an effector-to-memory phenotypical change after infusion.

### CAR T cells infiltrating the BM acquire a more differentiated and exhausted phenotype

As MM is usually characterize by the presence of malignant plasma cells in the BM and this may have an impact of CAR T cell phenotype, we next compared CAR T obtained from the BM or the PB and found only mild differences with an enrichment in CD8^+^ CAR T in both compartments with minor differences between patients (Figure 3A). Thus, we performed differential expression analysis between samples collected from BM and PB. Results showed 2142 differentially expressed genes (DEGs) among CD8^+^ CAR T cells, from which 1733 were upregulated in the BM and 409 in PB (|logFC>0.10| and FDR<0.05) (Table S5). Among genes upregulated in the BM we found markers related to activation and cytotoxicity (TNFRSF9/4-1BB, NKG7, PRF1, GZMH and GZMB), as well as terminal differentiation (DDX5, PRDM1) and exhaustion (LAG3, HAVCR2/TIM3). In contrast, genes expressed in PB were involved in memory (CD7), activation (FOS, JUN) and proliferation (TUBA1A), as supported by the statistical difference found in the G2M score among them (Figure 3B and S3A). We next focused on the analysis of DEG between BM and PB samples for each patient. Despite a limited number of shared genes between the 3 patients (Figure 3C and Table S6) we observed that genes upregulated in BM were enriched in pathways associated with T cell activation and cell adhesion (Figure 3D), providing common functional features of CAR T cells found in the BM.

**Figure 3.**
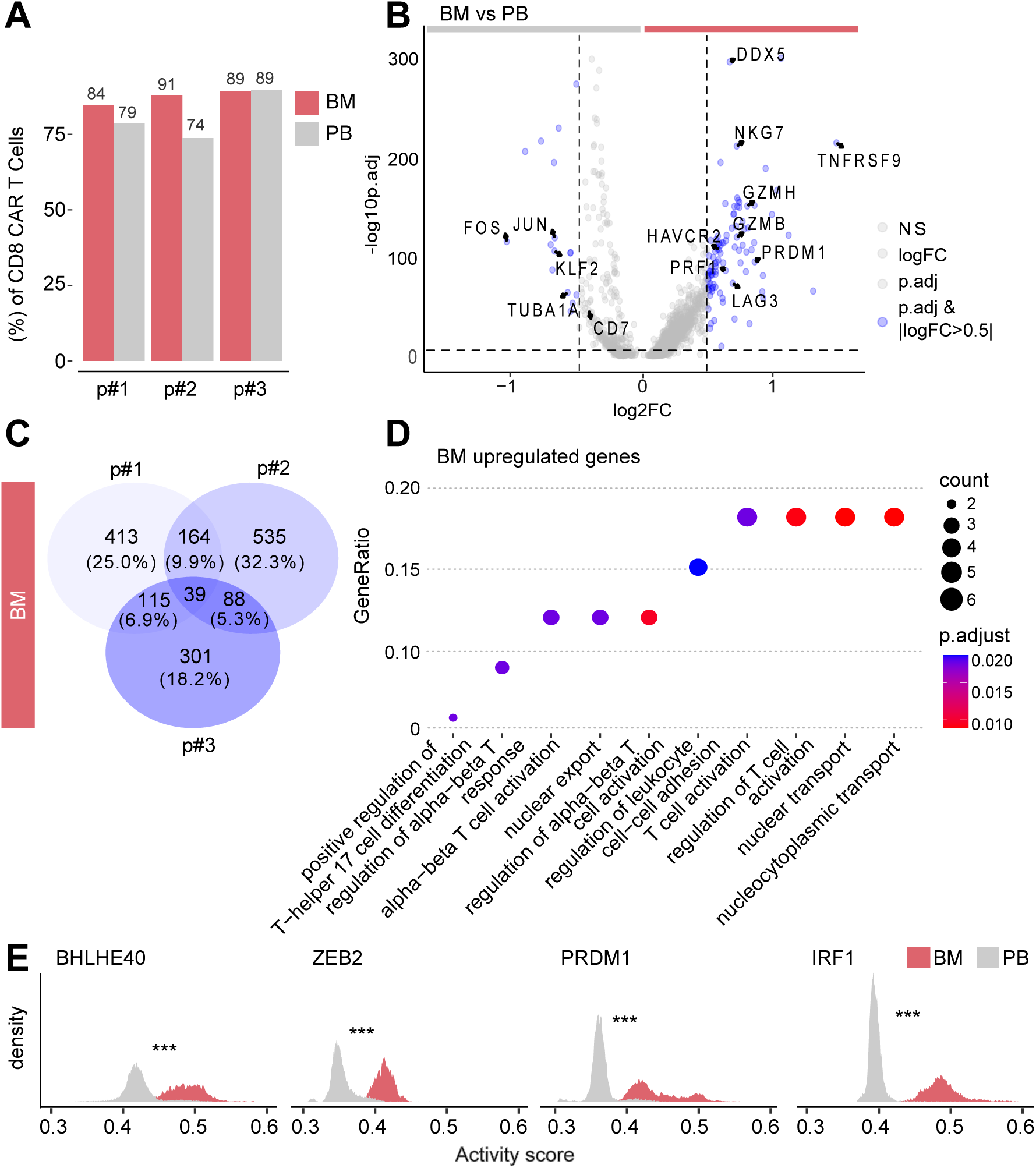
CAR T cells infiltrating the BM acquire a more differentiated and exhausted phenotype. Comparative analysis of paired CAR T cell samples from BM and PB was conducted. **(A)** Proportion of CD8^+^ CAR T cells per patient and location. **(B)** Differentially expressed genes between paired BM and PB samples (p adjusted < 0.01 and logFC > |0.5|). **(C)** Intersection of differentially expressed genes among BM samples of each patient. **(D)** Pathway enrichment analysis of genes upregulated in the BM in all three patients (p adjusted < 0.05). **(E)** Activity scores of BHLHE40, ZEB2, PRDM1 and IRF1 regulons in CAR T cells in BM and PB. Wilcoxon test for unpaired samples with Benjamini-Hochberg (BH) procedure for p value correction (B, E) and GO analysis with BH correction (D). ***p < 0.001.

GRN analysis revealed 99 regulons with differential activity between PB and BM (67 regulons more active in BM and 32 in PB) (Figure S3B and Table S7). Some of the regulons with increased activity in the BM included BHLHE40 and ZEB2, involved in T cell terminal differentiation^37,41^, PRDM1, associated to T cell dysfunction^34^, and IRF1, related to IFN response^42^ and antiproliferative responses against tumors (Figure 3E). On the other hand, TCF7, KLF2 and JUN family regulons were the most relevant regulons differentially active in PB (Figure S3C). The activation of this regulatory program is in consonance with the aforementioned general differences observed at transcriptomic level between PB and BM. All in all, these results suggest that in MM, the transcriptomic profile of post-infusion CAR T cells correlates with the presence of tumor cells which are predominantly in the BM. In that sense CAR T cells entering the BM acquire a terminal and cytotoxic phenotype leading to an overall more differentiated phenotype across the effector-to-memory axis in BM compared to PB.

### Limited effector-to-memory transition restricts CAR T cell persistence

Since our study included samples from three MM patients with different clinical characteristics in terms of CAR T cell persistence and therapeutic response (Table S8), we further analyzed patient-specific transcriptomic programs to understand the molecular mechanisms associated with different clinical behaviors. All three patients presented clear expansion of CAR T cells after infusion, with the peak of expansion at day 14 after infusion (described in^26,27^) CAR T cell expansion in the BM was observed in all three patients at day 30 post-infusion, albeit their levels differed among patients (Figure 4A). When delving into CAR T cell persistence, two of the patients presented almost undetectable CAR T cells from month 3 onwards in both PB and BM, while patient #3 presented long-term persistence with detectable CAR T cells up to month 23 after treatment (Figure 4A and Table S8). Interestingly, analysis of the distribution of the different CAR T cell populations revealed that the patient with long persistence (patient #3) presented a less differentiated profile in the effector-to-memory axis at early time points, in both BM and PB compartments, compared to those patients presenting low CAR T cell persistence (patients #1 and #2). Specifically, patient #3 had predominance of CD8^+^Eff in BM while in patients #1 and #2 there was a predominance of CD8^+^TE. Moreover, patient #3 presented predominantly CD8^+^EM cells in PB in comparison to the CD8^+^Eff observed in the other patients (Figure 4B).

**Figure 4.**
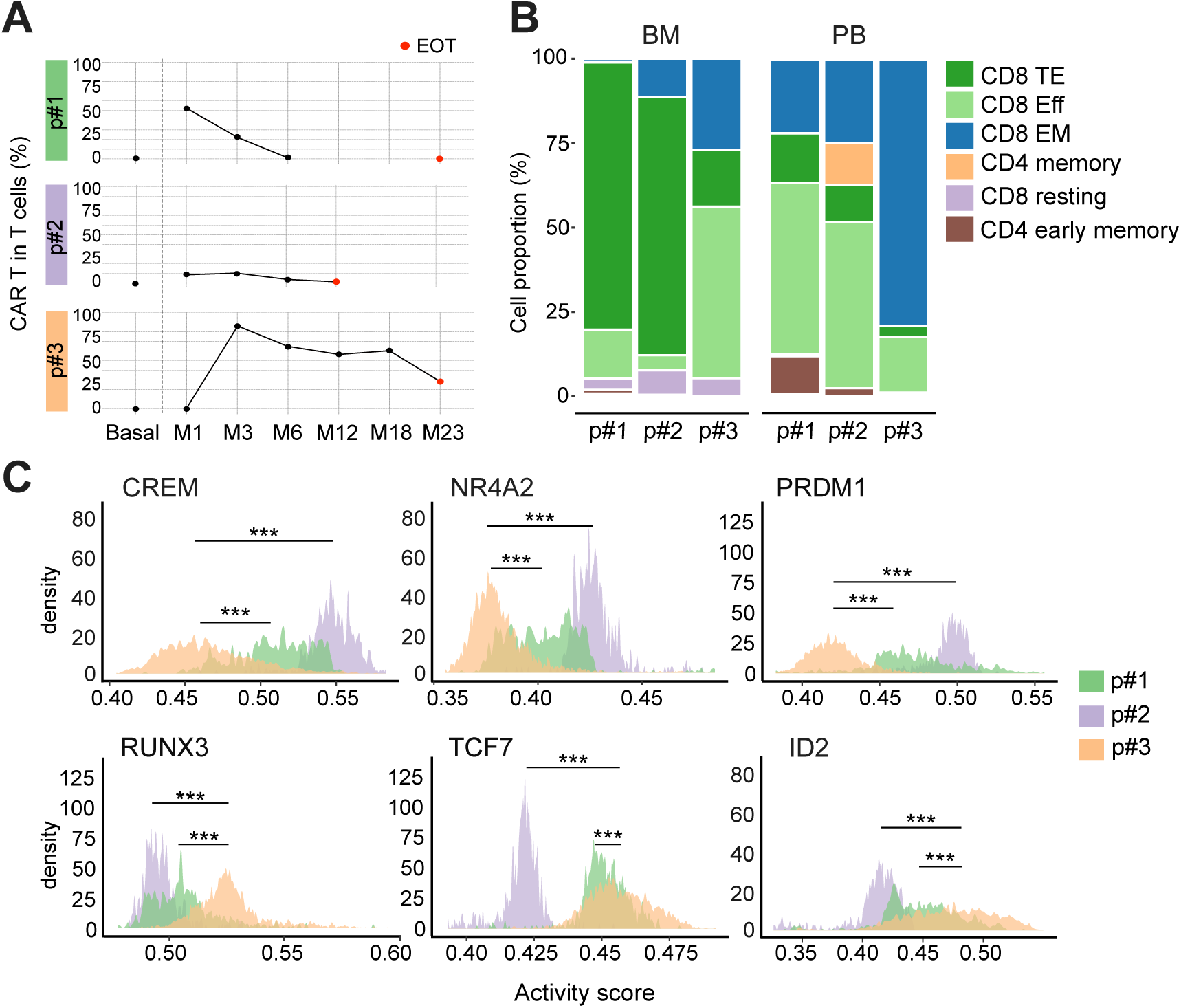
Gene regulatory dysfunctions in the effector-to-memory axis leads to lack of persistence. Transcriptomic features associated to long term CAR T cell persistence were analyzed. **(A)** Expansion of CAR T cells (% of T cells) in the BM in each patient. Data was obtained at different times after infusion until end of treatment (EOT). **(B)** Composition of post-infusion CAR T cells present in the BM and PB of each patient. **(C)** Activity scores of CREM, NR4A2, PRDM1, RUNX3, TCF7 and ID2 regulons in CAR T cells from BM of each patient. Kruskal-Wallis and Dunn’s test with Benjamini-Hochberg (BH) correction for post-hoc pairwise comparisons (C). ***p < 0.001.

Next, we explored GRNs involved in driving the observed differences across patients. As mentioned above, the specific regulatory program activated in CAR T cells infiltrating the BM presented a similar tendency across patients, supporting that there are conserved GRNs driving the CAR T cell activity. Nevertheless, the activity of some regulons varied notably across patients (Figure S4). Reduced activity of CREM, NR4A2 and PRDM1 regulons, involved in T cell dysfunction and exhaustion, along with increased activity of RUNX3, TCF7 and ID2, important factors for T cell infiltration^43^ and memory formation, was observed in the patient with the longest CAR T cell persistence (Figure 4C). This would be in accordance with the differential distribution of the effector/effector memory CAR T cell populations observed between patients. In summary, the differential activity of factors related to CAR T cell dysfunction like PRDM1 and NR4A2, (increased in patients #1 and #2), along with the higher activity of factors related to memory (TCF7 and ID2) in patient #3, might explain the differences in CAR T cell persistence, suggesting that the activity levels of some regulons could be key to determine the fate of CAR T cells *in vivo*.

### Clonality analysis identifies a hyperexpanded clone in the BM of a patient with partial response

Regarding the antitumoral response, 90% of the patients enrolled in CARTBCMA-HCB-01 clinical trial presented a very good partial response or better during the first 100 days from infusion^26,27^, while 7% had a partial response. Interestingly, patient #1 and #3 had a stringent complete response, achieving MRD negativity at month 1. In contrast, patient #2 had a partial response with disease progression in month 11 after treatment (Table S8). No transcriptional differences between patients in PR and in sCR were observed. The analysis of scTCR-seq revealed that CAR T cell diversity was similar among most samples (Figure 5A), with a general decrease in clonotype repertoire after infusion (Figure S5A), according to what has already been described^18^, but maintaining >75% of CAR T cells with a clonotype detected only once (unique clonotypes) (Figure 5B). Interestingly, CAR T cells from BM of the patient with partial response (patient #2) presented lower values in Shannon and ACE scores (Figure 5A), and a reduced clonotype repertoire with only around 30% of unique clonotypes (Figure 5B and S5A). In fact, most of the clonal space of that sample was occupied by a single clone (herein referred to as clonotype 1), that comprised around 70% of all the sequenced CAR T cells (Figure 5C). Clonal CAR T cell expansions, although not very frequent, have been observed in previous studies analyzing BCMA targeting CAR T cells^25,44^. The hyperexpanded clone mostly belonged to the aforementioned CD8^+^TE cell population (Figure 5D), and it was not found in its paired PB sample nor in the IP, suggesting a specific clonal expansion within the BM (Figure 5C).

**Figure 5.**
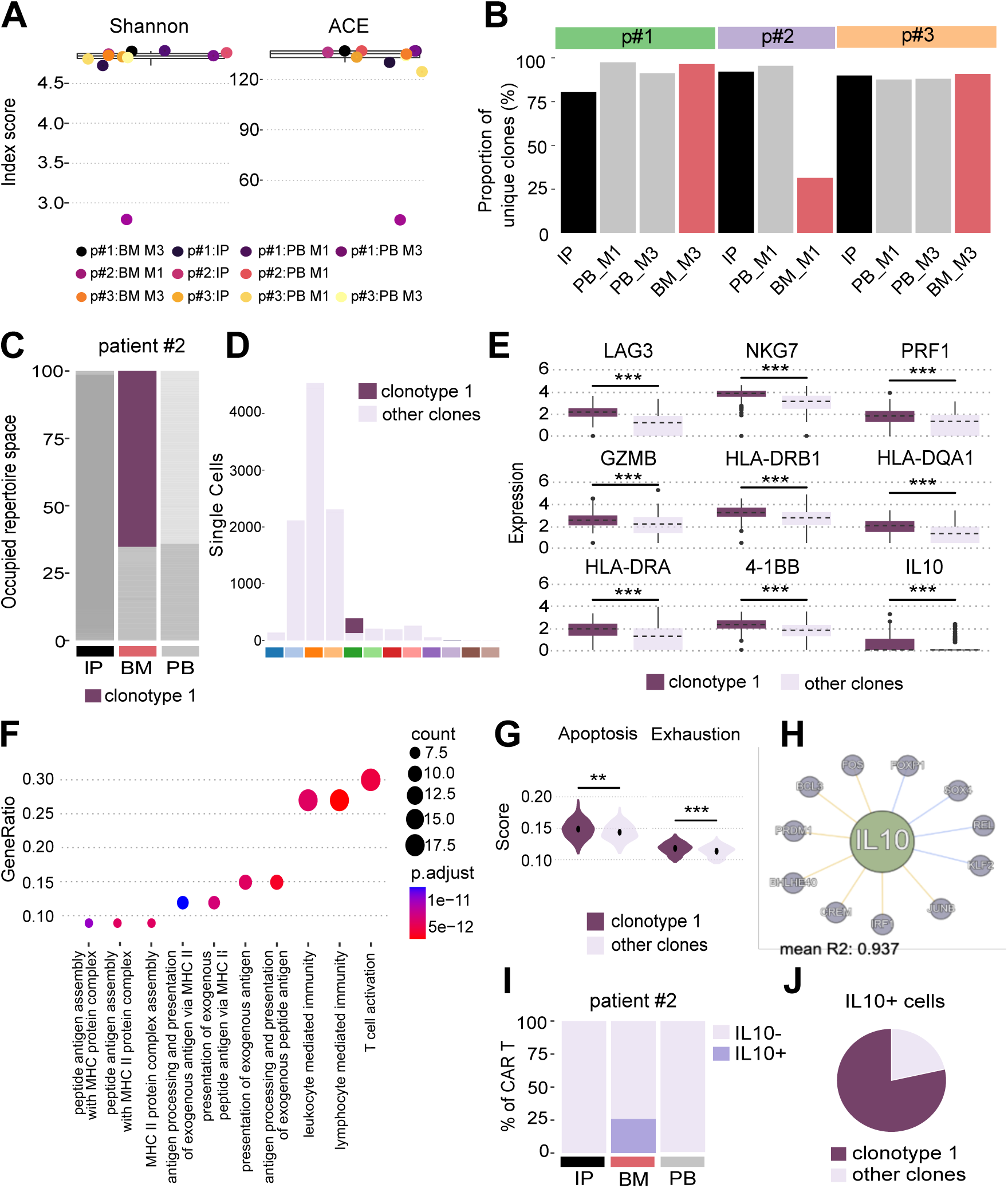
Clonality analysis reveals a hyperexpanded clone in the BM of a patient with partial response. Clonal repertoire of CAR T cells from the different samples was evaluated by scTCR-seq. **(A)** Shannon and Abundance-based coverage estimator (ACE) scores of each sample. **(B)** Percent of unique clonotypes in each sample. **(C)** Relative space occupied by the hyperexpanded clonotype 1 in the different sample from patient #2. **(D)** Distribution of the hyperexpanded clonotype 1 within the different CAR T cell populations. **(E)** Differential expression of the indicated genes in clonotype 1 compared to other clones. **(F)** Analysis of pathway enrichment with genes upregulated in the clonotype 1 compared to other clones (p adjusted < 0.05). **(G)** Apoptosis and Exhaustion signature scores between clonotype 1 and other clones. **(H)** IL10 regulon depicting IL10 as target and the TFs potentially regulating its activity indicating their positive (yellow) or negative (blue) correlation. **(I)** Percent of IL10-expressing cells within the CAR T cells from IP, BM and PB samples of patient #2. **(J)** Percent of clonotype 1 cells among IL10-expressing cells in the BM sample of patient #2. Wilcoxon test for unpaired samples (E, G) and GO analysis (F). Benjamini-Hochberg (BH) procedure for p value correction (E, F). **p < 0.01, and ***p < 0.001.

The analysis of transcriptomic differences between clonotype 1 and the other clonotypes present in the same sample showed upregulation of genes related activation (HLA-DQA1, HLA-DRA, HLA-DRB1, TNFRSF9/4-1BB), cytotoxicity (NKG7, PRF1, GZMB) as well as exhaustion (LAG3) (Figure 5E and S5B). Moreover, GO analysis of DEGs in clonotype 1 revealed pathways related to T cell activation, antigen processing and lymphocyte mediated immunity (Figure 5F). In addition, clonotype 1 was enriched in genetic signatures associated to apoptosis and exhaustion (Figure 5G). Interestingly, we observed that IL10, a molecule associated with antiproliferative and immunosuppressive functions, was also upregulated in clonotype 1 (Figure 5E and S5B). IL10 has been shown to be produced by highly activated effector CD8^+^ cells at the peak of infection^45,46^. Accordingly, a regulatory network analysis with IL10 as target gene, revealed TFs associated to effector functions and terminal differentiation, like FOS, BHLHE40, PRDM1 and CREM, as potential regulators of IL10 production (Figure 5H), suggesting that IL10 production in clonotype 1 could be a consequence of its highly cytotoxic phenotype. A closer look showed that IL10^+^ cells were only present in BM and were mostly expressed by clonotype 1 cells (Figure 5I and 5J). Further analysis of IL10 producing cells in other patients showed that they were very scarce in the BM of patient 1 and undetectable in patient 3 (Figure S5C). To expand the analysis to a bigger number of patients, we looked for cells expressing IL10 in a publicly available single cell data set of post-infusion BCMA CAR T cells^25^, however we did not find IL10^+^ cells among post-infusion samples (Figure S5D). We also searched for IL10 in two different single cell data sets of CD19 CAR T cells collected at short (< 1year) and long (1 to 6 years) post-infusion^17,18^. IL10^+^ CAR T cells were only found in the two patients from the long-term study (Figure S5E and S5F), and despite the presence of CD8^+^ CAR T cells after infusion, most of IL10^+^ cells were CD4^+^ or double negative, and not cytotoxic CD8^+^ cells as found in our data (Figure S5G and S5H). Altogether, these results suggest that clonal expansion of IL10 expressing CD8^+^ CAR T cells might be associated with reduced clinical efficacy.

### TCR stimulation strengthens CAR T cell activation and promotes IL10 production

CAR T cell activation usually leads to polyclonal or oligoclonal cell proliferation, and clonal expansions have been proposed to be a consequence of additional factors, including TET2 mutation or CAR T hyperactivation^18,22,44^. In this sense, we further evaluated whether endogenous TCR engagement may promote clonal CAR T cell expansion leading to the specific transcriptomic characteristics of the identified clonotype 1. First, we observed that CAR T cells from the BM of patient #2 were enriched in a TCR-specific signature generated for previously published data^29^ (see materials and methods) compared to their PB counterparts (Figure 6A and Table S9) suggesting the activation of the endogenous TCR. Similarly, BM IL10^+^ cells also presented a higher score compared to BM IL10^-^ cells, supporting the idea that TCR stimulation could lead to IL10 production (Figure 6A). To further explore the role of the TCR and the CAR receptors in IL10 production, we generated CAR T cell from OT-1 mice, allowing us a specific stimulation of the endogenous TCR and/or the CAR (Figure 6B). We observed a dose-dependent production of IL10 and IFN-γ upon TCR stimulation with SIINFEKL peptide, as well as upon CAR stimulation with mouse recombinant BCMA (rBCMA), confirming CAR T cell activation and cytokine release through both receptors (Figure S6A and S6B). When TCR and CAR were simultaneously stimulated an additive effect was observed in IL10 production compared to the stimulation of either receptor alone (Figure 6C), in accordance with the cooperation that had been formerly described^47^. Interestingly, we also observed a similar additive effect when the receptors were stimulated sequentially (CAR stimulation for 48h followed by the TCR) (Figure 6C), suggesting that CAR-activated T cells were able to produce increased levels of IL10 upon TCR engagement. Next, we studied the effect that IL10 could exert on the functionality of the surrounding CAR T cells by adding recombinant IL10 (rIL10) to the culture media. Our results showed a significant antiproliferative effect of IL10 in stimulated CD8^+^ CAR T cells, together with increased IFN-γ production (Figure 6D and S6C), which may in turn lead to CAR T cell exhaustion through expression of immune checkpoint inhibitors^48^.

**Figure 6.**
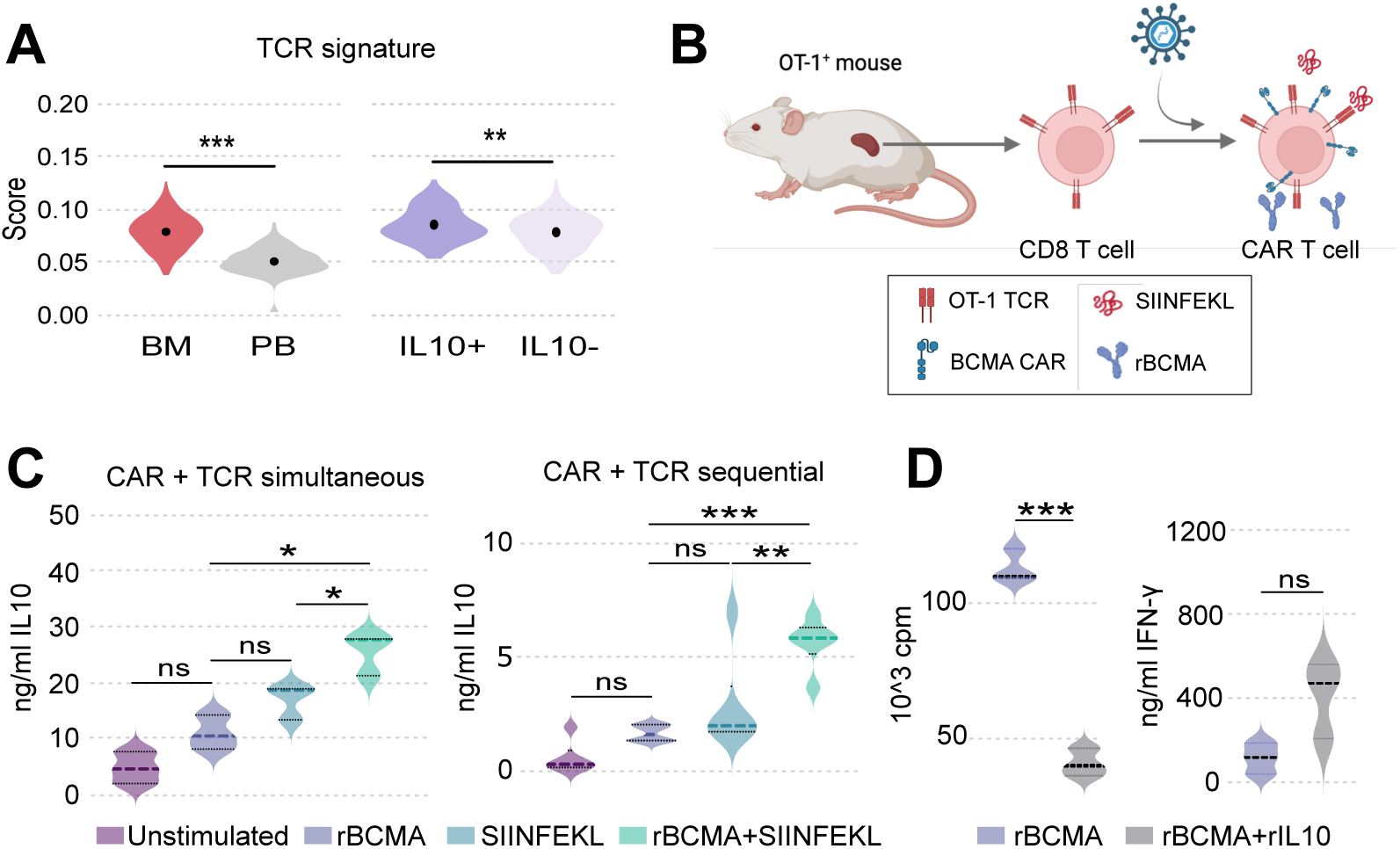
TCR stimulation strengthens CAR T cell activation and promotes IL10 production. The effect of TCR stimulation on CAR T cell proliferation and functionality was evaluated. **(A)** TCR signature score of the CAR T cells from BM and PB samples (left), and of the IL10^+^ and IL10^-^ cells (right) of patient #2. **(B)** Schematic representation of the generation of BCMA CAR T cells from OT1^+^ mice. **(C)** Quantification of IL10 levels (ng/ml) in supernatants of mouse OT1^+^ CAR T cells stimulated with SIINFKEL, rBCMA, or a combination of both simultaneously (left) or in a sequential manner (right). **(D)** Quantification of OT1^+^ CAR T cell proliferation (left) or IFN-γ levels in supernatants of OT1^+^ CAR T cells (right) after incubation during 48h with recombinant IL10. Wilcoxon test for unpaired samples (A), one-way ANOVA with Turkey test for p value correction in post-hoc pairwise comparisons (C) and student’s t-test for unpaired samples with Welch’s correction (D, E). ns p>0.05, *p < 0.05, **p < 0.01, and ***p < 0.001.

Overall, our data show that CAR T cell activation through CAR and TCR engagement increases IL10 production, and that produced IL10 modifies the behavior of CAR T cells reducing cell proliferation and promoting IFN-γ production. Thus, these results provide a potential mechanism by which infections and/or pathogens may alter CAR T cell functionality once they have been infused into the patients. Moreover, these results highlight the need to take into consideration the potential contribution of the TCR during CAR T therapy, to have a full vision of the mechanisms governing CAR T cell function.

## DISCUSSION

Chimeric antigen receptor (CAR) T cell therapy has transformed the treatment landscape for relapsed/refractory multiple myeloma (MM) improving both OS and DFS. However, challenges remain as relapses of the disease remains constant. In this study, we leveraged single-cell multiomics to dissect molecular and transcriptional mechanisms implicated in CAR T cell efficacy and persistence in MM patients treated with BCMA-CAR T cells. Our results showed a preferential expansion of CD8^+^ CAR T cells 1 and 3 months after infusion both in the BM and the PB, whereas CD4^+^ CAR T cells were underrepresented. However, the low expression levels of the CD4 gene may compromise accurate cell annotation in single-cell data, meaning that some double-negative (CD4⁻CD8⁻) cells could actually be CD4⁺ CAR T cells, leading to an underestimation of this population. Nonetheless, even in this scenario the number of CD4^+^ CAR T cells would remain low, without a significant impact on the observed preferential expansion of CD8⁺ CAR T cells. This expansion is in line with previous studies in patients treated with BCMA CAR T cells where CD8⁺ CAR T cells predominantly expand in the early post-infusion period^23,25^. Moreover, while the presence of long-lasting CD4^+^ CAR T cells has been associated with long term remission in patients with leukemia^17^, the preferential expansion of CD8^+^ CAR T cells was also observed in some of those patients at earlier times^17^, suggesting the importance of an initial cytotoxic CD8^+^ T cell response for proper CAR T cell antitumoral functionality.

The analysis of paired samples demonstrated that CAR T cells infiltrating the BM acquire a more activated, yet terminally exhausted phenotype compared to their PB counterparts. This was evidenced by the increased expression of exhaustion markers (e.g., LAG3, HAVCR2) and the activation of key transcription factors such as PRDM1 and ZEB2, which have been associated with terminal differentiation and T cell dysfunction^37,49,50^. Conversely, PB CAR T cells exhibited a higher expression of memory-related markers (TCF7^51^ and JUN^20,35^) and regulatory networks supporting persistence. Interestingly, overexpression of JUN in CAR T cells has been shown to reduce terminal differentiation in various mouse tumor models^52^, aligning with our observation that PB provides a more favorable environment for maintaining a functional CAR T cell pool. These findings support the concept that the interaction of the MM plasma cells with the CAR T cells in the BM led to their terminal differentiation and that the PB may constitute a transient reservoir of CAR T cells with a more persistent phenotype. Our results are consistent with reports from Rade et al. that found BCMA CAR T cells to be more activated in the BM compared to PB^25^.

Clonality analysis provided additional insights into CAR T cell dynamics among patients. Patients who achieved complete response (CR) exhibited greater TCR diversity across all samples analyzed. In contrast, the patient who did not achieve CR exhibited a more homogeneous TCR repertoire in the BM, driven by the dominance of the IL10-producing clone showing upregulation of activation and cytotoxicity markers. This clone, belonging to the CD8 terminal effector cluster, was not detected in the IP or paired PB sample. Clonal expansion in CAR T cells from MM patients at one-month post-infusion has been previously described^25^ as a common event in BCMA CAR T therapy. Interestingly, while previous reports observed greater clonal expansion in CR patients compared to non-CR patients^25^, and expansion in both PB and BM, we found hyperexpansion only in the BM of a patient with partial response (PR), suggesting that clonal proliferation occurred within the BM^53^. Though traditionally associated with immunoregulatory functions of CD4^+^ T cells^54^, IL10 has been shown to be produced by effector CD8^+^ T cells during peak infection, likely as a means of reducing inflammation^45^. Moreover, several studies have challenged the view of IL10 as solely immunosuppressive^55,56^, demonstrating its ability to induce tumor-specific responses of CD8^+^ T cells and enhance immune surveillance in mouse models^57^. A recent study reported beneficial effects of IL10-secreting CAR T cells in solid tumors^58^. Using various mouse tumor models, the authors showed enhanced granzyme B and IFN-γ production by IL10-producing CAR T cells compared to conventional CAR T cells^58^. In line with these findings, we observed increased levels of IFN-γ in the culture media of CAR T cells incubated with recombinant IL10. Additionally, we found that the elevated levels of IFN-γ were, at least in part, due to an increased number of cells producing the cytokine.

Given that IL10 was predominantly expressed by the expanded clone, we hypothesized that the same factors responsible for clonal expansion could also drive IL10 production. By generating OT-1^+^ CAR T cells, we demonstrated that IL10 production occurred upon engagement of both CAR and TCR receptors in a dose-dependent manner. At low antigen levels, CAR and TCR stimulation resulted in additive IL10 production, even when the receptors were not simultaneously activated. This suggests that IL10 production may not only be a result of strong CAR or TCR activation but also of weaker, sustained engagement of both receptors. Therefore, the elevated IL10 levels observed in the expanded clone may reflect either strong transient activation or prolonged contact with lower antigen levels. Our findings support the hypothesis that TCR engagement may contribute to CAR T dysfunction in MM raising the intriguing possibility that interactions between the CAR and endogenous TCR may play an underappreciated role in shaping CAR T cell fate as has been previously described^59,60^.

From a clinical perspective, our findings provide actionable insights for optimizing CAR T cell therapies in MM. First, engineering CAR T cells with enhanced memory-like properties (e.g., by overexpressing TCF7 or inhibiting PRDM1) may improve persistence. Second, strategies to mitigate excessive exhaustion in BM-infiltrating CAR T cells, such as checkpoint blockade or metabolic modulation, should be explored. Finally, further investigation into the role of IL10 and TCR engagement in CAR T dysfunction could pave the way for novel interventions to prevent clonal exhaustion and improve response durability. In summary, our study, combining scRNA-seq/scTCR-seq allowed the identification of key molecular mechanisms governing post-infusion BCMA CAR T cell responses providing key transcriptional regulators, clonotypic dynamics, and potential immunosuppressive factors such as IL10 promoting CAR T cell dysfunction, that would represent a potential target to be modulated for the development of improved CAR T therapies for MM.

## Acknowledgments

We thank the members of Hematology and Cell Therapy Department of the Clinica Universidad de Navarra for input throughout the course of the project and all the patients as well as families who made this study possible. We particularly acknowledge the patients for their participation in the Clinical trial CARTBCMA-HCB-01 (NCT04309981) and the Biobank of the University of Navarra for its collaboration.

## Author contributions

M.H., J.R.R.-M., and F.P. designed the study. L.J-U., G.S., S.C. and M.E.C-C. performed the data analysis. L.J-U., T.L., J.J.L. and J.R.R-M. designed experimental validations. L.J-U., S.R-D., E.I. and T.L. performed wet lab experiments. P.R-O., A.A-P., E.T. and J.S-M. provided clinical advice. A.Z., A.O-C., M.E-R., V.C., J.L.R., V.G-C., M.V.M., F.S-G., S.I., A.L-D., A.G-N., M.J., C.F.d.L., and B.P. provided clinical samples and/or data. P.S.M-U. performed single cell library preparation and sequencing. P.S.M-U. and D.A. provided technical assistance. L.J-U., G.S., S.C., M.H., J.R.R.-M., and F.P. discussed the study design and the results. M.H., J.R.R.-M., and F.P. were responsible for research supervision, coordination and strategy. L.J-U. and J.R.R.-M. drafted the manuscript. M.H., J.R.R.-M., and F.P. reviewed and edited the manuscript. All authors reviewed and approved the final version of the manuscript.

## Data and materials availability

All data needed to evaluate the conclusions in the paper are present in the paper and/or the Supplemental Materials. The scRNA-seq and scTCR-seq data generated in this study have been deposited in the GEO database (GSE290061).

