## Supplemental_Material for "Single-Cell Multiomics Reveals Regulatory Mechanisms of CAR T Cell Persistence and Dysfunction in Multiple Myeloma"

### Extended Material and Methods

**FACS-sorting of CAR T cells from patient PBMCs and BM.**
CAR T cells were FACS-sorted from Bone Marrow (BM) aspirates and Peripheral Blood (PB) samples. BM and PB samples were bulk lysed to remove erythrocytes, and they were surface-marked in a two-step procedure with an antibody targeting the scFv of the CAR (ScFv-BT, Jackson Immunoresearch #115-065-072) and Streptavidine-BV421 (BD #563259). Specifically, 2.5µl of ScFv-BT were added in 100µl FACS buffer per sample tube, and they were incubated for 30 at RT in the dark. Then, samples were washed 3x times with 4ml PBS containing 0.04% BSA and Strep-BV421 was added (1µl in 100µl). Then, CAR T cells were isolated using either BD FACSAria Ilu or Moflo ASTRIOS BC.

**5' Gene expression Profiling5' Gene expression Profiling.**
Single-cell RNA sequencing (scRNA-seq) was conducted using the Chromium Single Cell 5′ Reagent Kit (10X Genomics), following the manufacturer's guidelines. In brief, CAR T cells were sorted into 100 µl of 1X PBS containing 0.04% BSA, and their concentration and viability were assessed using a Nexcelom Cellometer K2 Fluorescent Cell Counter. Cell concentration was adjusted to a range of 700–1200 cells/µl to optimize the recovery target. The single-cell suspension was then combined with RT Master Mix and co-loaded with barcoded 5′ single-cell gel beads and partitioning oil into separate single-cell A Chips to create Gel Beads in Emulsion (GEMs) using the Chromium Controller. Inside each GEM, cells were lysed, polyadenylated RNA was reverse transcribed, and the resulting cDNA was barcoded. Post-GEM RT cleanup was used for recovery of the barcoded complementary DNA (cDNA), and followed by PCR amplification. The cDNA was quantified using the Qubit dsDNA HS Assay Kit (Thermo Fisher Scientific), and its quality was evaluated using the High Sensitivity D5000 ScreenTape Assay (Agilent Technologies). A total of 50 ng of double-stranded cDNA was used for constructing the 5′ Gene Expression library. The cDNA was fragmented, end-repaired, A-tailed, and ligated to an adaptor. A final round of PCR amplification with barcoded primers was performed for sample indexing. Sequencing was carried out on a NextSeq500 (Illumina) with Read1: 26 cycles, Read2: 57 cycles, and i7 index: 8 cycles, achieving an average depth of 30,000 reads per cell.

**Single-cell TCR repertoire sequencing.**
TCR α/β sequencing was performed using the 10X Genomics Single Cell V(D)J Immune Profiling Solution (10X Genomics). Briefly, full-length V, D, and J gene segments were amplified from barcoded cDNA using the Chromium Single Cell V(D)J Enrichment Kit (for human T cells) via two consecutive PCRs with primers targeting both the adaptor and the gene constant region. The enriched cDNA was quantified as described above, and library construction was carried out as described above with 50 ng cDNA. After quality control and quantification, sequencing of the Single Cell V(D)J enriched libraries was outsourced to Macrogen Inc. (http://www.macrogen.com/) with a minimum sequencing depth of 5,000 reads per cell.

**Raw sequencing data processing, QC, data filtering, and normalization.**
Raw scRNA-seq data was demultiplexed, aligned to the human reference (GRCh38) and the feature-barcode matrix was quantified using Cell Ranger (v6.0.1) from 10X Genomics. Further computational analysis was performed using Seurat (v3.1.5). Cells were filtered based on several quality control (QC) parameters, including the number of detected genes, number of UMIs, and the proportion of UMIs mapping to mitochondrial and ribosomal genes per cell. Thresholds for each parameter were determined by examining their distribution within each single-cell library and through visual inspection of QC scatter plots. In general, cells with less than 250 genes and cells with less than 500 UMIs were discarded. Cells with more than 30000 UMIs were also discarded to remove likely doublet or multiplet captures. Genes present in fewer than three cells were also excluded from further analysis, as well as low-quality cells, including those with over 10% mitochondrial content. Finally, cells with no detectable expression of ARI002h (gene encoding for the CAR molecule) were discarded to avoid potential contamination from CAR negative T cells and other mononuclear cells.

**Unsupervised cell clustering and dimensionality reduction.**
All datasets were merged and log-normalized, and highly variable genes were identified while removing unwanted sources of variation using Seurat (v3.1.5). Harmony (v0.1.0) was used for integration of all datasets. Different resolution parameters were tested for unsupervised clustering to determine the optimal number of clusters. In this study, the first 20 principal components (PCs) and 2000 highly variable genes identified by Seurat were used for sample integration and clustering, with a resolution

parameter set at 0.8, resulting in the identification of 12 cell clusters. Dimensionality reduction was performed using t-distributed stochastic neighbor embedding (t-SNE) and Uniform Manifold Approximation and Projection (UMAP).

**Inferring cell cycle stage and DEGs.**

The cell cycle stage of individual cells was computationally assigned using Seurat's R code, which is based on the expression patterns of cell-cycle-related signature genes. Differentially expressed genes (DEGs) for each cluster were identified using the Seurat FindConservedMarkers or FindAllMarkers function with the Wilcoxon Rank Sum Test and Benjamini-Holchberg (BH) procedure for p value correction.

**Determination of major cell types and cell states.**

The primary cell types (CD4 and CD8) were defined by the expression of marker genes (CD4, CD8A, and CD8B) according to the 10X Genomics transcriptome data. To further classify the cell types and states (such as activated, effector or memory), a manual review of DEGs for each cell cluster was conducted, referencing canonical marker genes.

**TCR V(D)J sequence assembly, paired clonotype calling, TCR diversity, and clonality analysis.**

TCR reconstruction and paired TCR clonotype analysis were conducted using Cell Ranger (v6.0.2) for V(D)J sequence assembly.

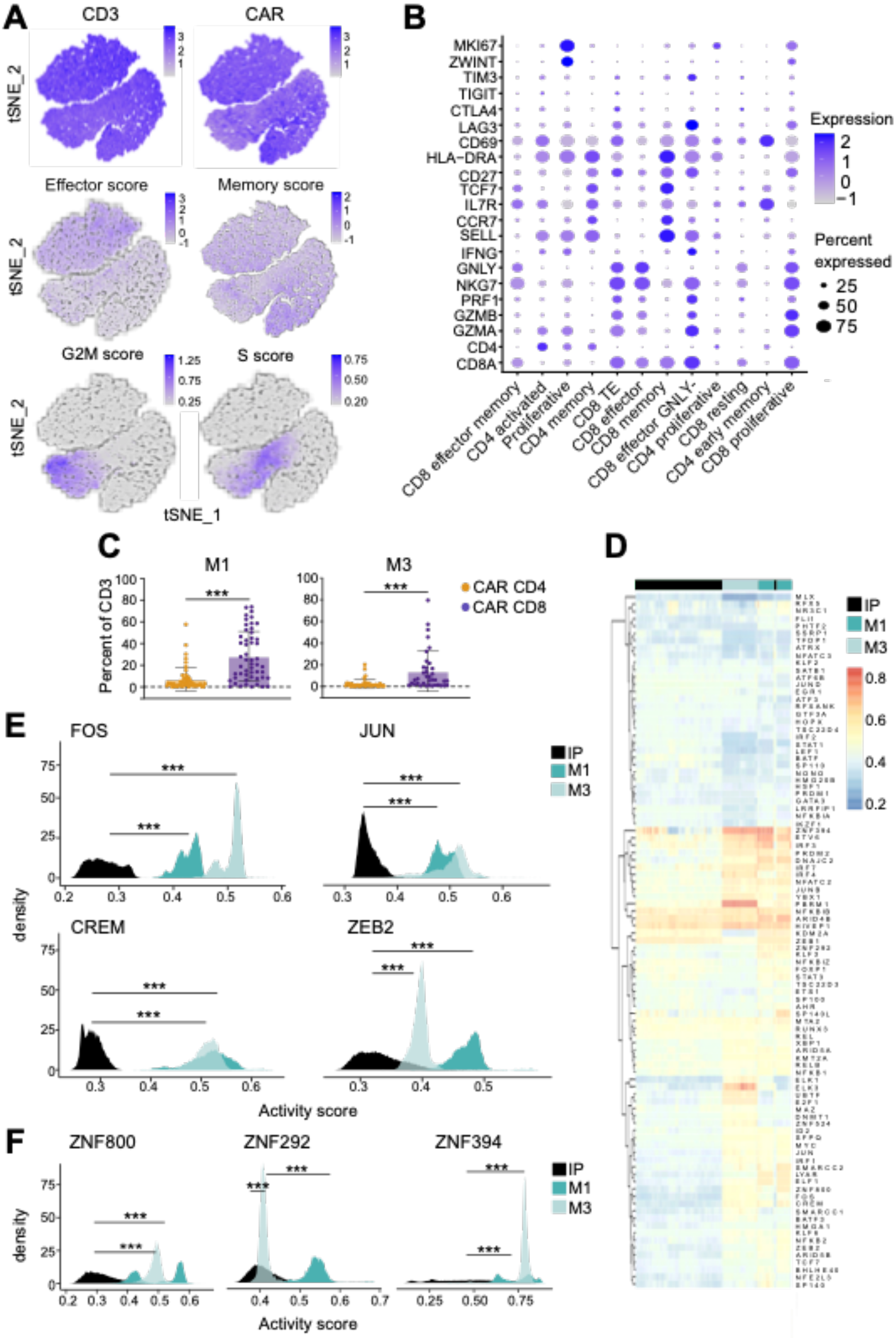


**Supplemental Figure 1. Characterization of CAR T cells by scRNA-seq. (A)** t-SNE representation of CAR T cells showing CD3E and CAR expression after quality control. **(B)** Dot plot representing the average expression of canonical T cell markers in each cluster. **(C)** Percentage of $CD4^{+}$ and $CD8^{+}$ CAR T cells at 1 and 3 months after infusion in the BM of patients enrolled in CARTBCMA-HCB-01 trial. **(D)** Heatmap showing the activity of regulons in IP and post-infusion samples as inferred by SimiC. **(E)** Density plots depicting the activity scores of selected regulons with increased activity after infusion. **(F)** Density plots depicting the activity of selected regulons composed by zinc finger factors with increased activity after infusion. Wilcoxon test for paired samples (C) and Kruskal-Wallis test and Dunn's test with Benjamini-Hochberg (BH) correction for post-hoc pairwise comparisons (D, F). ***p < 0.001.

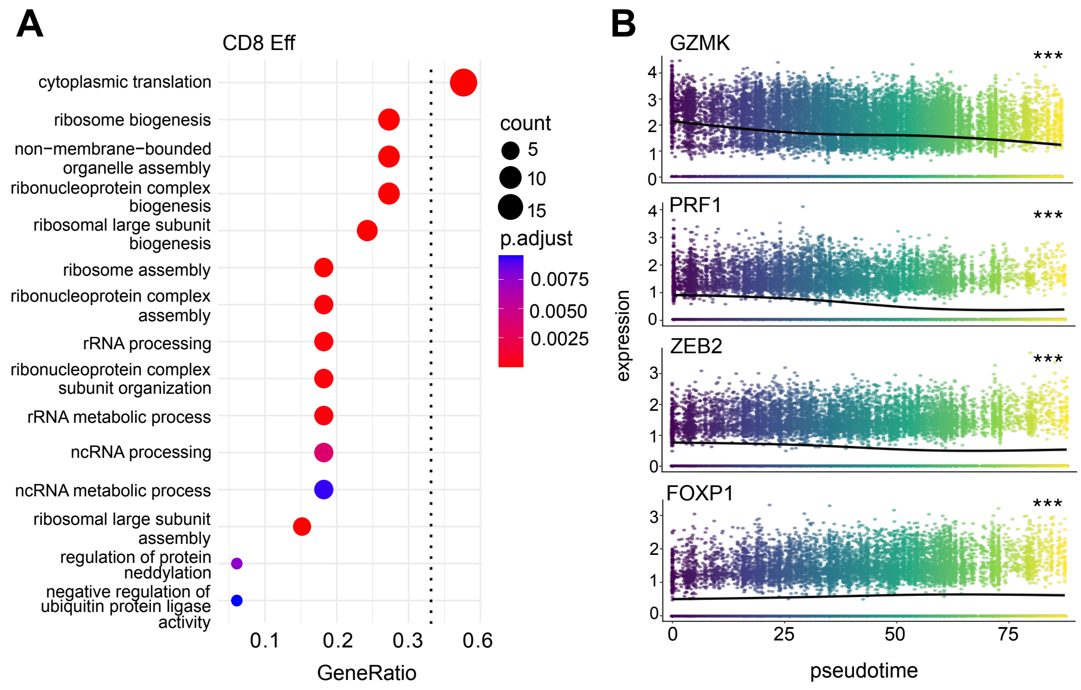


**Supplemental Figure 2. Transcriptional differences among main post-infusion populations. (A)** GO analysis of genes upregulated in $CD8^+$ effector cluster when compared to $CD8^+$ terminally-differentiated effector cells (p adjusted < 0.05). **(B)** Expression of GZMK, PRF1, ZEB2 and FOXP1 along the effector-memory axis. GO analysis and Benjamini-Hochberg (BH) procedure for p value correction (A) and natural cubic splines regression (B). ***p < 0.001.

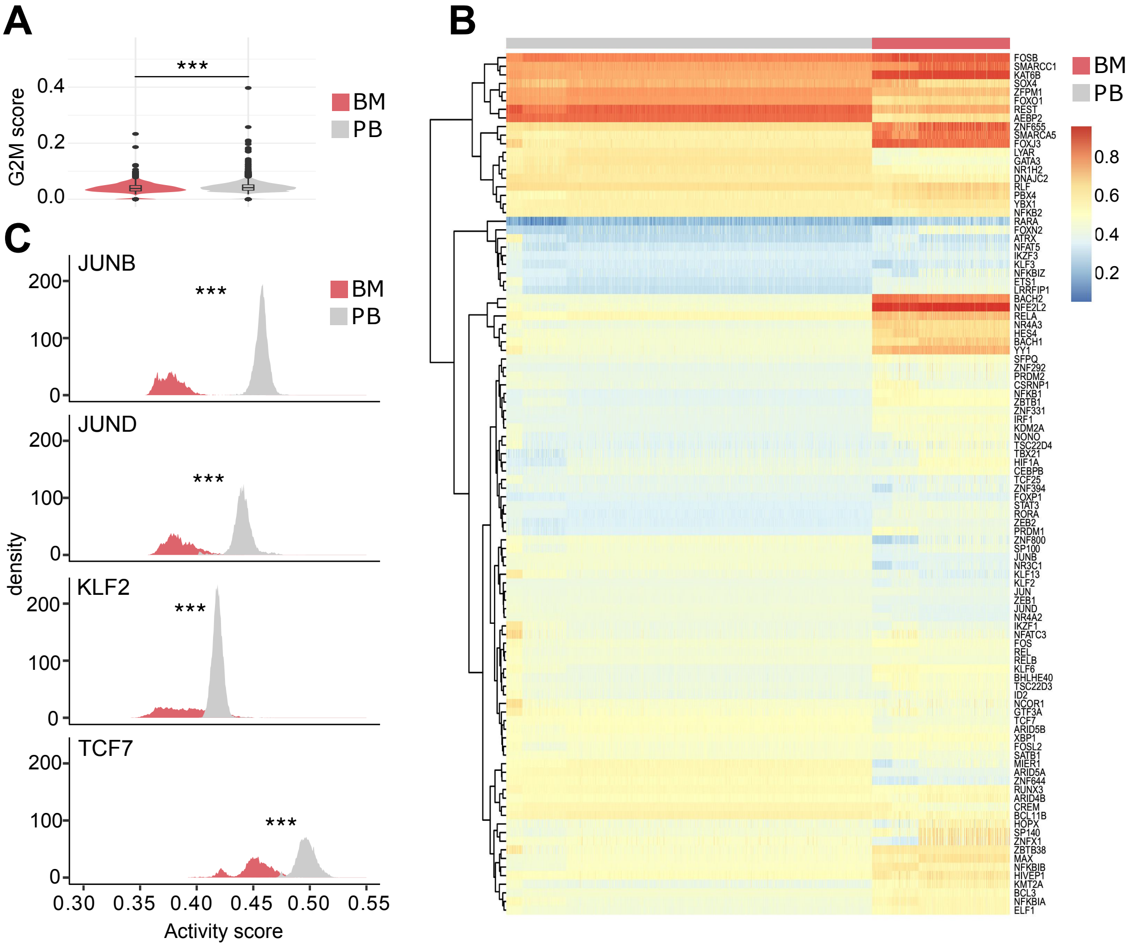


**Supplemental Figure 3. Transcriptional differences between BM and PB CAR T cells. (A)** G2M score values in BM and PB CAR T cells as scored by UCell. **(B)** Heatmap showing the activity scores of regulons in BM and PB CAR T cells as computed by SimiC. **(C)** Density plots reflecting the activity scores of JUNB, JUND, KLF2 and TCF7. Wilcoxon test for unpaired samples (A, C) and Benjamini-Hochberg (BH) procedure for p value correction (C). ***$p < 0.001$.

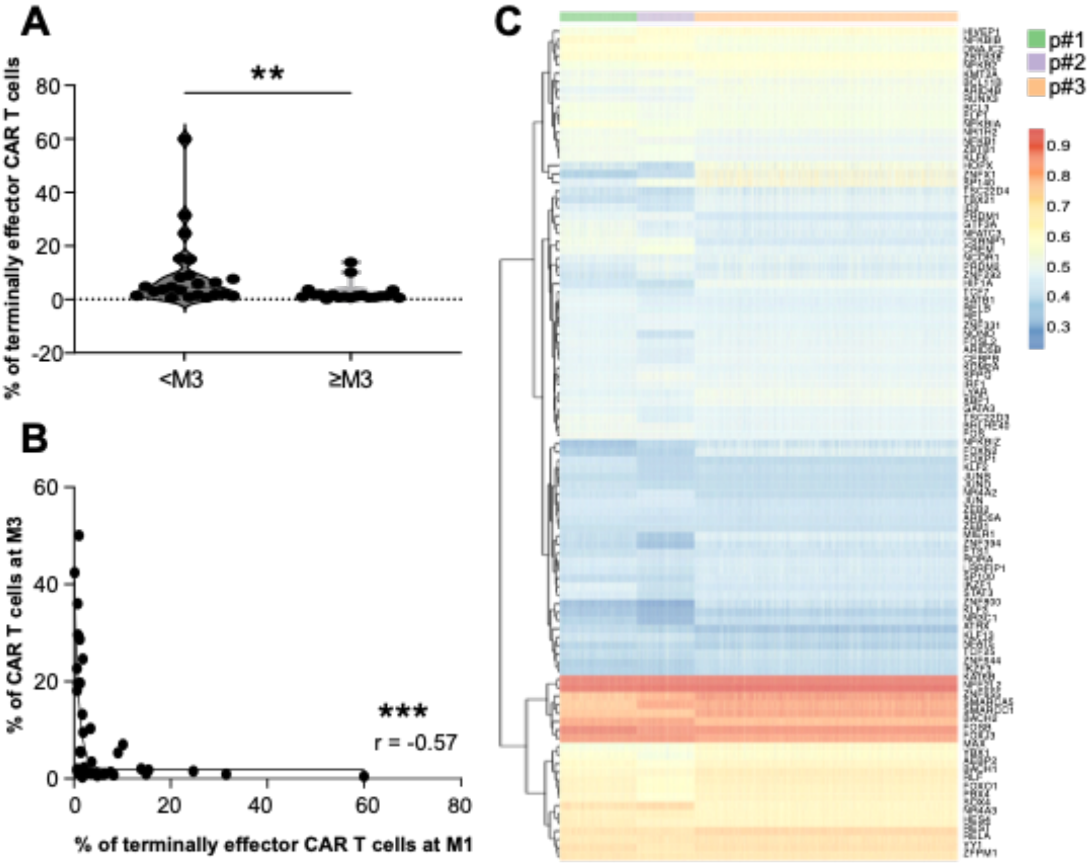


**Supplemental Figure 4. Differences in BM GRN across patients.**

**A)** Phenotypic analysis of CAR T cells present at month 1 in the BM of all the patients included in CARTBCMA-HCB-01 clinical trial. Percentage of terminally effector cells ($CD62L^{-}/CD45RA^{+}$) present in patients with low (<M3) and long (≥M3) CAR T cell persistence. **B)** Correlation analysis between the levels of terminally effector CAR T cells at month 1 and the expansion of CAR T cells (percent of T cells) at month 3. **C)** Heatmap showing differences across patients in the activity scores of indicated regulons in the CAR T cells present at the BM. Mann Whitney test for unpaired samples (A) and Spearman correlation analysis (B). **$p < 0.01$; ***$p<0.001$.

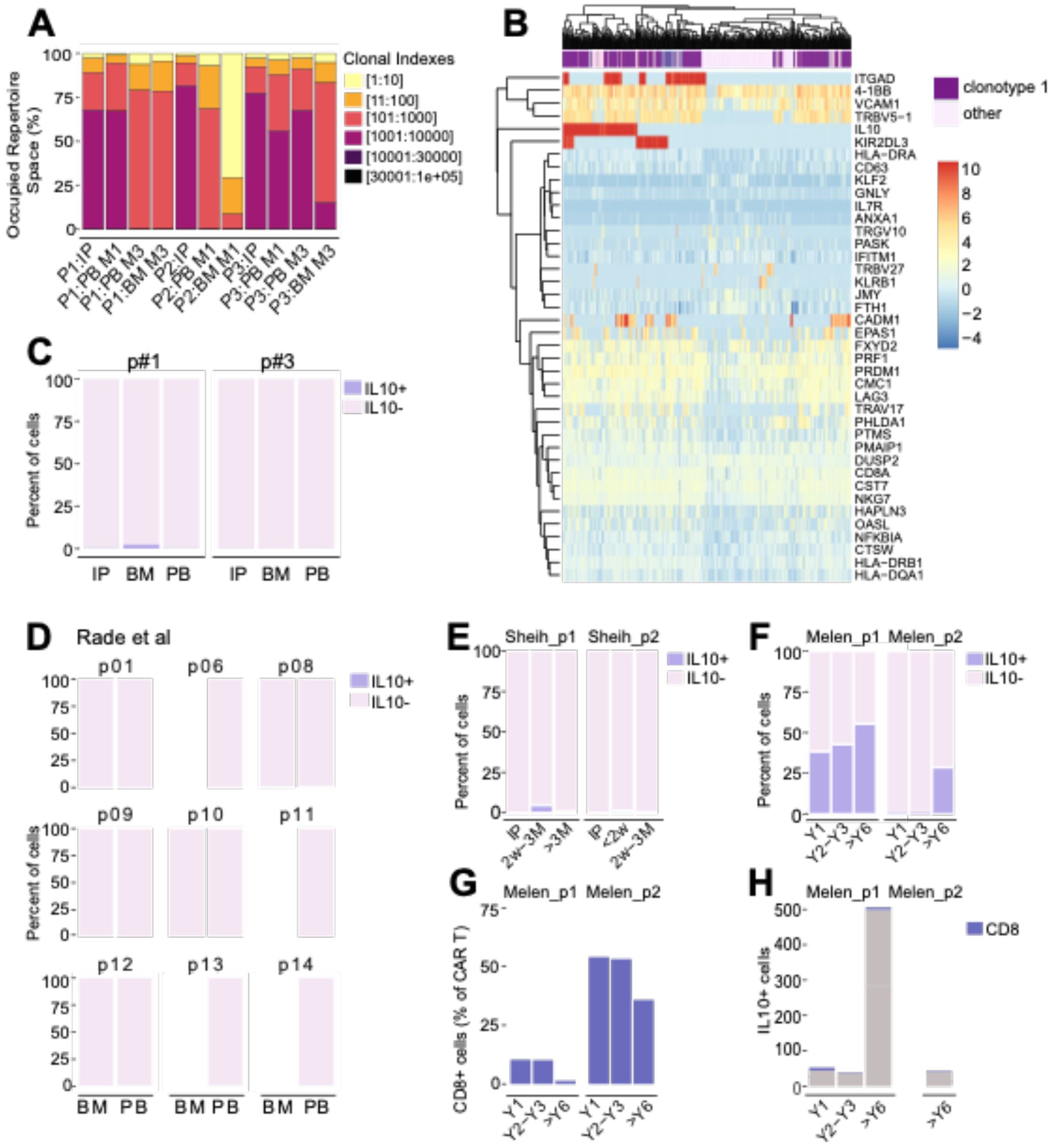


**Supplemental Figure 5**. **Analysis of clonotypes. (A)** Proportion of clonal repertoire in each sample occupied by clonotypes grouped by relative abundance. **(B)** Heatmap depicting the expression of DEGs between clonotype 1 and other clonotypes in the same sample (p adjusted <0.05). **(C)** Percent of cells expressing IL10 (=> 1 count) in patients #1 and #3 per sample. **(D)** Percent of cells expressing IL10 (=> 1 count) in a publicly available T single cell data set after BCMA CAR T cell infusion. **(E)** Percent of cells expressing IL10 (=> 1 count) in a publicly available single cell data set of CD19 CAR T cells sequenced at short times after infusion (< 1 year). **(F)** Percent of cells expressing IL10 (=> 1 count) in a publicly available single cell data set of CD19 CAR T cells sequenced at long times after infusion (1 to 6 years). **(G)** Proportion of $CD8^+$ CAR T cells in two patients' data from the long-term CD19 CAR T single cell data set. **(H)** Cell counts of $CD8^+$ cells among $IL10^+$ cells in the two patients of the long-term CD19 CAR T single cell data set.

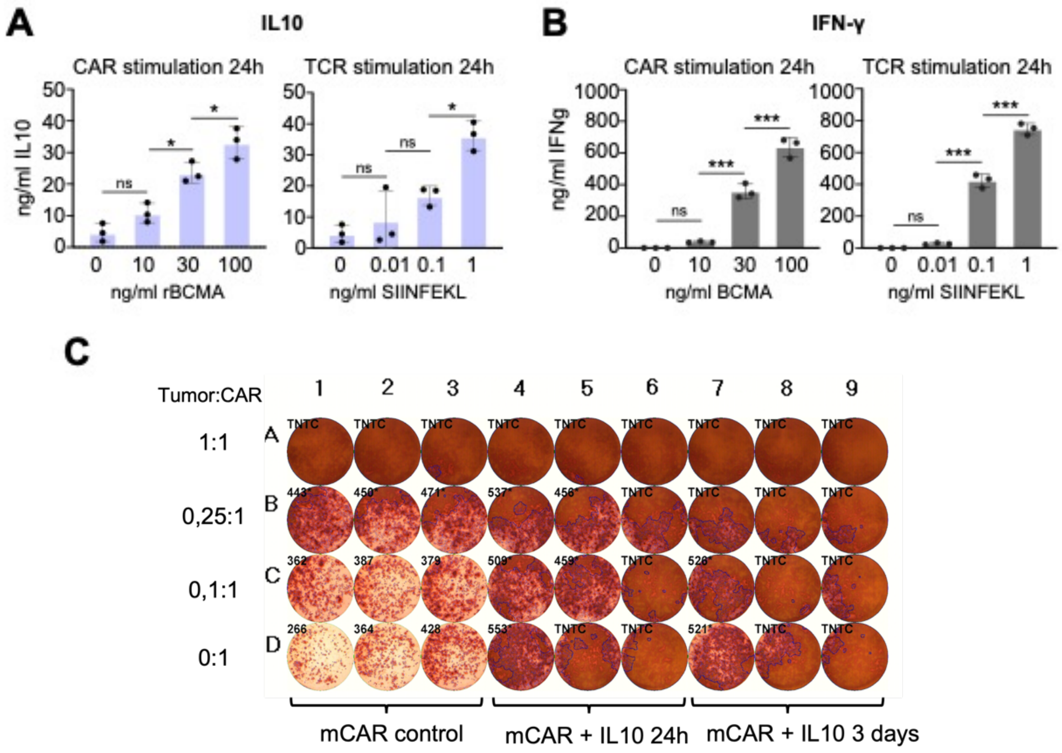


**Supplemental Figure 6. *In vitro* exploration of TCR and CAR stimulation. (A)** Barplots showing the production of IL10 upon 24h CAR (left) or TCR (right) stimulation with rBCMA and SIINFEKL, respectively. **(B)** Barplots showing the production of IFN-ɣ upon 24h CAR (left) or TCR (right) stimulation with rBCMA and SIINFEKL, respectively. **(C)** Picture of IFNγ ELISPOT of mouse CD8 OT1$^+$ CAR T cells stimulated with recombinant IL10 for 24h and 72h. One-way ANOVA with Turkey test for p value correction in post-hoc pairwise comparisons (A, B). ns $p>0.05$, *$p < 0.05$ and ***$p < 0.001$.

### Supplementary Tables

**Table S1. Number of analyzed CAR T cells per sample.**

| | Peripheral Blood | | | Bone Marrow | |
|---|---|---|---|---|---|
| **Patient** | **IP** | **M1** | **M3** | **M1** | **M3** |
| **#1** | 8655 | 4444 | 1036 | | 628 |
| **#2** | 10282 | 384 | | 469 | |
| **#3** | 8781 | 3847 | 7189 | | 2140 |

**Table S8. Clinical characteristics of the patients included in the study.**

| | Sex | Age | CR at M12 | MRD positivity | CAR T persistence in BM | CRS | Degree of CRS | ICANS | Infections |
|---|---|---|---|---|---|---|---|---|---|
| **#1** | Male | 67 | Yes | N/A | 6 months | Yes | 2 | No | |
| **#2** | Female | 62 | No | M11 | 12 months | Yes | 1 | No | At M10 |
| **#3** | Male | 53 | Yes | M18 | >18 months | Yes | 2 | No | Respiratory infection - d11<br>CMV reactivation - d31 |

**Table S2. List of differentially expressed genes between IP and post infusion samples.**
Attached as a separate file.

**Table S3. Activity of regulons in IP vs post-infusion samples.**
Attached as a separate file.

**Table S4. List of differentially expressed genes between post-infusion effector populations.**
Attached as a separate file.

**Table S5. List of differentially expressed genes between BM vs PB CAR T cells.**
Attached as a separate file.

**Table S6. List of common differentially expressed genes across patients.**
Attached as a separate file.

**Table S7. Activity of regulons in BM vs PB samples.**
Attached as a separate file.

**Table S9. List of the genes used for the generation of the signatures.**
Attached as a separate file.
